# Metabotropic group II glutamate receptors mediate cue-triggered increases in incentive motivation for reward

**DOI:** 10.1101/2021.09.14.460377

**Authors:** Caroline Garceau, Justine Marsault, Mike J F Robinson, Anne-Noël Samaha

**Affiliations:** Department of Pharmacology and Physiology, Faculty of Medicine, Université de Montréal, Montréal, QC, H3T 1J4, Canada; Department of Neurosciences, Faculty of Medicine, Université de Montréal, Montréal, QC, H3T 1J4, Canada; CNS Research Group, Faculty of Medicine, Université de Montréal, Montréal, QC, H3T 1J4, Canada

**Keywords:** Instrumental conditioning, Pavlovian conditioning, Pavlovian-to-instrumental transfer, glutamate, mGlu2/3 receptors, devaluation, sex differences

## Abstract

**Rationale:** Reward-associated cues can acquire incentive motivational properties and invigorate reward-seeking actions via Pavlovian-to-instrumental transfer (PIT). Glutamatergic neurotransmission mediates the appetitive effects of reward-associated cues. We characterized the expression of PIT and its mediation by metabotropic group II glutamate (mGlu2/3) receptor activity in female and male rats.

**Objectives:** Across the sexes, we used PIT procedures to determine i) cue-triggered increases in incentive motivation for water reward (Experiment 1), ii) the respective influences of the mGlu2/3 receptor agonist LY379268 and reward devaluation by satiation on this effect (Experiment 2).

**Methods:** Water-restricted male and female Sprague-Dawley rats learned to lever press for water. Separately, they learned that one of two auditory stimuli predicts free water (CS+ vs CS-). On PIT test days, the CS+ and CS- were presented non-contingently, and we measured effects on lever pressing under extinction (no water). In Experiment 1, we characterized PIT across the sexes. In Experiment 2, we measured PIT after systemic LY379268 administration (0, 0.3 and 1 mg/kg), and water satiation, respectively.

**Results:** Female and male rats showed similar PIT, with CS+ but not CS- presentations potentiating water-seeking behaviour. LY379268 (1 mg/kg) attenuated CS+ evoked increases in both water-associated lever pressing and conditioned approach to the water port. Reward devaluation attenuated both water-seeking and CS+ evoked conditioned approach behaviour.

**Conclusions:** The sexes show similar cue-triggered increases in reward ‘wanting’, and water devaluation suppresses both water seeking and cue-triggered anticipation of water reward. Finally, across the sexes, mGlu2/3 receptor activity mediates cue-triggered increases in reward ‘wanting’.

## Introduction

Sights, sounds and smells in the environment that signal the occurrence of rewards can acquire incentive motivational properties, enabling such cues to trigger ‘wanting’ for both the cues and the rewards they predict. This motivational state in turn guides animals towards rewards such as food, water, shelter, and safety. Control over behaviour by reward cues is required for survival. For example, when animals are hungry or thirsty, they must be responsive to environmental cues signaling food or water. In parallel, when reward cues are attributed with too little or too much incentive motivational value, this can contribute to disorders such as depression and addiction, respectively. Thus, understanding the neurobiological mechanisms underlying cue-triggered incentive motivation is relevant to both adaptive and maladaptive reward-seeking behaviours.

Glutamatergic neurotransmission mediates the behavioral response to reward-associated cues (Di Ciano, Cardinal, Cowell, Little, & Everitt, 2001), and growing evidence suggests that this involves activity at mGlu2/3 receptors. These receptors are mainly presynaptic and are coupled to Gi signaling, such that their stimulation decreases synaptic glutamate release (Conn & Pin, 1997; Imre, 2007; Schoepp, 2001). Injecting the mGlu2/3 receptor agonist LY379268 either into the central nucleus of the amygdala (Lu, Uejima, Gray, Bossert, & Shaham, 2007; Uejima, Bossert, Poles, & Lu, 2007), into the nucleus accumbens core (Bossert, Gray, Lu, & Shaham, 2006) or systemically (Backstrom & Hyytia, 2005; Baptista, Martin-Fardon, & Weiss, 2004; Bossert, Poles, Sheffler-Collins, & Ghitza, 2006; Zhao et al., 2006) suppresses cue-induced reinstatement of reward-seeking behaviour. These studies highlight the contributions of mGlu2/3 receptors to reward seeking, but cue-induced reinstatement procedures are not pure cue-triggered incentive motivation tests. Indeed, cue-induced reinstatement procedures also involve secondary reinforcement, as instrumental responding at test is reinforced by cue presentation (Shaham, Shalev, Lu, de Wit, & Stewart, 2003).

In contrast, Pavlovian-to-Instrumental transfer or (PIT) is a pure conditioned incentive paradigm that measures the ability of a cue to increase motivation for a reward (Cartoni, Balleine, & Baldassarre, 2016; Rescorla & Solomon, 1967; Walker, 1942; Wyvell & Berridge, 2000). In PIT, subjects learn to perform an instrumental response for a primary reward. Separately, they learn that one Pavlovian cue predicts non-contingent delivery of that reward (CS+), and a second, distinct cue does not (CS-). On the PIT test day, the rats can perform the same instrumental response, but no reward is given, and both the CS+ and CS- are presented non-contingently throughout this test. PIT is seen when CS+ presentations trigger increases in ongoing instrumental responding, indicating cue-triggered increases in incentive motivation for the reward. PIT avoids primary reinforcement effects, because subjects are tested under extinction conditions, so that no primary reward is delivered. PIT also avoids secondary reinforcement, because instrumental responding at test is not reinforced by the reward cue.

Using PIT, we found previously that injecting the mGlu2/3 receptor agonist LY379268 into the BLA suppressed cue-triggered increases in ‘wanting’ for the associated water reward (Garceau, Samaha, Cordahi, Servonnet, & Khoo, 2021). This work identifies potential neural circuits within which activation of these receptors mediates cue-triggered increases in reward ‘wanting’. However, unlike systemic drug administration, intracerebral administration leads to the inevitable question of translational potential to humans.

In addition, to our knowledge, no published study [including Garceau et al. (2021)] has reported on the contributions of mGlu2/3 receptor activity to cue-triggered increases in incentive motivation in female animals. There are limitations to generalizing findings obtained with males to females (Becker & Koob, 2016; Prendergast, Onishi, & Zucker, 2014; Shansky, 2019; Shansky & Murphy, 2021), and there are both similarities and differences in how the sexes respond to reward cues. Studies comparing the sexes on cue reactivity report mixed results, with some studies showing more (Perkins et al., 2001; Robbins, Ehrman, Childress, & O’Brien, 1999), less (Sterling, Dean, Weinstein, Murphy, & Gottheil, 2004) or comparable (Avants, Margolin, Kosten, & Cooney, 1995; Negrete & Emil, 1992; Rubonis et al., 1994; Waldrop et al., 2010) effects of drug-associated cues in drug-using women relative to men. Female and male rats can also show both similarities and differences in how they respond to appetitive cues, as measured using cue-induced reinstatement of cocaine-seeking behaviour (Feltenstein, Henderson, & See, 2011) or Pavlovian extinction of PIT (Delamater, Schneider, & Derman, 2017).

Thus, here, we compared female and male rats on both cue-triggered increases in incentive motivation for a water reward, and the influence of systemic administration of a mGlu2/3 receptor agonist on this effect.

## Methods

### Animals

Adult male (200-225 g) and female (150-175 g) Sprague-Dawley rats (Charles River Laboratories, Montréal, Quebec, Canada) were housed 2 per cage in a climate-controlled (22±1°C, 30±10% humidity) colony room, under a reverse 12-hour light/dark cycle (lights off at 8:30 a.m.). We trained and tested rats during the dark phase of the circadian cycle. Upon arrival, food (Rodent 5075, Charles River Laboratories) and water were freely available in the home cage. Rats were handled daily after 72 hours of acclimation to the animal colony. Since we used water as the unconditioned stimulus (US), rats had restricted access to water beginning at 4 days after their arrival to facilitate acquisition of Pavlovian and Instrumental conditioning. Rats initially had 6 h/day of access to water for the first 4 days, 4 h/day for the next 3 days, and then 2 h/day until the end of each experiment. We gave water at least 1 hour after the end of testing and removed water bottles at the same time each day. Daily water consumption during each two-hour water restriction period was measured throughout Experiment 2. The Université de Montréal approved all procedures involving rats and procedures followed the ‘Principles of laboratory animal care’ and the guidelines of the Canadian Council on Animal Care.

### Behavioural Apparatus

Rats were trained and tested in standard operant chambers (31.8 x 25.4 x 26.7 cm; Med Associates, VT, USA) located in a testing room separate from the rats’ housing room. The operant chambers were placed within light and sound-attenuating boxes equipped with a ventilation fan that masked external noise. Each chamber was equipped with a tone generator located adjacent to the house light on the back wall, and a clicker located outside the chamber. On the opposite wall, two retractable levers (left active; right inactive) were located on either side of a water cup equipped with an infrared head entry detector. A liquid dispenser was set to deliver 100-μL drops of water into the cup. Each chamber also contained 4 infrared photobeams spaced evenly at floor level to measure locomotor activity. A computer running Med-PC IV was connected to chambers and collected data.

#### Exp. 1: Cue-triggered increases in incentive motivation for water reward in female and male rats

##### Instrumental conditioning

Behavioral procedures for general PIT were adapted from Derman and Ferrario (2018) and Garceau et al. (2021). Figure 1a shows the training and testing timeline. Female and male rats (n = 16/sex) first underwent instrumental conditioning, during which they were trained to lever press for water reward. In the first session, water (100 μL) was delivered on a fixed-ratio 1 reinforcement schedule (FR1). Pressing a second lever (inactive) produced neither the reward nor any other programmed consequence. Sessions ended after 40 min or when rats earned 50 water deliveries. Rats were required to earn 50 water deliveries within a session to transition from FR1 to a variable interval (VI) reinforcement schedule. We increased the VI schedule across 8 sessions (40 min/session), in the following order: two VI10 sessions (range: 5-15 sec), two VI30 sessions (range: 15-45 sec) and four VI60 sessions (range: 30-90 sec). To determine the extent to which rats learned the response-water association, we recorded lever pressing and number of water rewards earned throughout each training session.

**Fig. 1.**
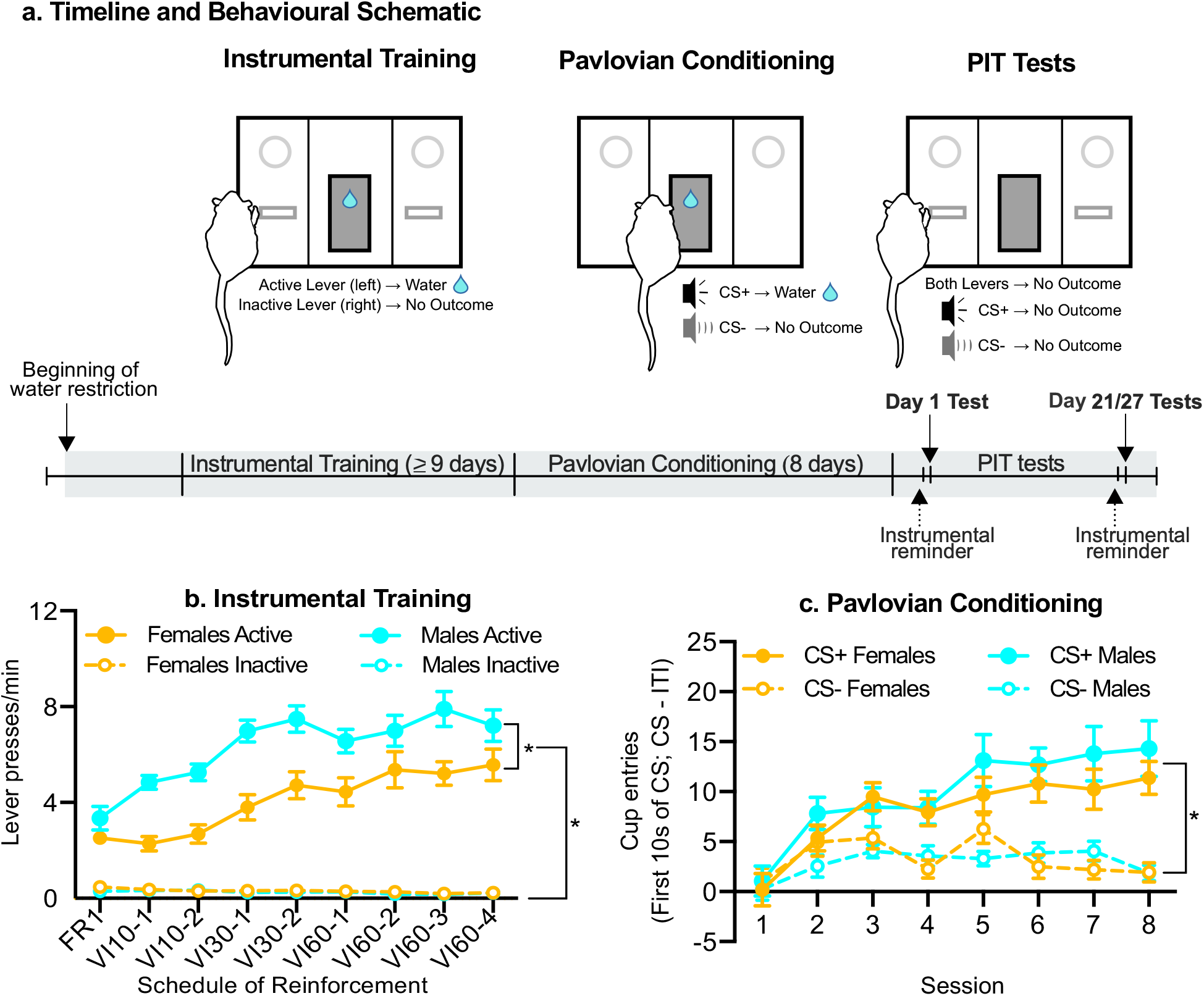
Timeline and acquisition of instrumental and Pavlovian conditioning in Experiment 1. (a) Female and male rats received instrumental (lever pressing for water reward) and Pavlovian (CS+/water and CS-/no water conditioning) sessions. We then measured cue-triggered increases in incentive motivation for water reward using Pavlovian-to-instrumental transfer (PIT) procedures. (b) Rats rapidly learned to discriminate between active and inactive levers, and the rate of active-lever presses increased over the daily 40-min instrumental training sessions, with the males pressing more often on the active lever than females did. (c) The rate of water cup entries during the first 10 sec of each CS presentation increased over the daily 44-min Pavlovian conditioning sessions, and there were no sex differences in this response. Data are means ± SEM (n = 16/sex). * *p* < 0.05. FR; fixed ratio. VI; variable interval. CS; conditioned stimulus. ITI; inter-trial interval.

##### Pavlovian conditioning

Next, rats underwent 8 Pavlovian conditioning sessions during which they received eight, 2-minute presentations of two distinct auditory stimuli (4 presentations each); either a 1800-Hz, 85-dB tone or a 10-Hz clicker. One stimulus (CS+) was paired with 4 water deliveries (100 μL each) per CS presentation on a VI30 schedule (range: 15-45 s; first water delivery ≥ 10 s from CS onset), while the other stimulus (CS-) was presented an equal number of times, but had no programmed consequences. Rats were allocated to the tone-CS+ or clicker-CS+ conditions such that mean active lever presses across the last four VI60 sessions and the number of FR1 sessions required to earn 50 water deliveries/session were comparable across the two CS+ conditions. On average, each intertrial interval (ITI) separating CS presentations lasted 180 s (range: 120-240 s), such that each Pavlovian session lasted 44 min. The CS+ and CS- were presented in alternation during each session. During Pavlovian conditioning, levers were always retracted, such that rats could not interact with them. To determine the extent to which rats learned the CS+-water association, we recorded water cup entries during CS presentations and during the ITI.

##### Pavlovian-to-Instrumental transfer (PIT) testing

To compare the sexes on the expression and persistence of cue-triggered increases in incentive motivation for water reward, rats were tested for PIT following Pavlovian conditioning (‘Day 1’) and then again three to four weeks later (‘Days 21-27’). On the day prior to each PIT test, rats received an instrumental reminder session identical to VI60 training described above. Levers were available throughout each PIT test session (42 min). PIT tests were conducted under extinction conditions, such that lever pressing never produced water. This enables assessment of cue-triggered increases in incentive motivation for the primary reward, without the confounding influence of primary reinforcement. After an initial 10 minutes, each CS (CS+ and CS-) was presented 4 times (2 min/presentation) in a counterbalanced order, with a fixed 2-min ITI. The CS were presented non-contingently, and never in response to lever pressing. This avoids the confounding influence of secondary reinforcement. We measured lever responses, water cup entries and locomotion throughout the session.

#### Exp. 2: Effects of systemic injection of an mGlu2/3 receptor agonist on cue-triggered increases in incentive motivation for water reward

Figure 3a illustrates the experimental timeline for Experiment 2. A new cohort of male and female rats (n = 16/sex) were water restricted, trained, and tested as described for Experiment 1. PIT testing began two weeks after the end of Pavlovian conditioning.

##### LY379268 administration

Rats received a subcutaneous injection of saline, 0.3 or 1 mg/kg of the mGlu2/3 receptor agonist LY379268 (Cat No.: 2453, Batch No.: 9B/232416, CAS No.: 191471-52-0, Tocris Bioscience, Oakville, Ontario; in a volume of 1 ml/kg), 30 min before the start of PIT testing. We used a within-subjects design, with doses given in counterbalanced order (1 dose/day). LY379268 was dissolved in 0.9% saline, and the solution was then briefly heated and sonicated. The pH was adjusted to ~7 with ~0.5 μl of 10N NaOH per 1 mg of LY379268 (Allain, Roberts, Levesque, & Samaha, 2017; Imre et al., 2006). Between PIT tests, rats received 1 instrumental and 1 Pavlovian reminder session.

##### Effects of reinforcer devaluation on cue-triggered incentive motivation

In Experiment 1, female and male rats were compared on the expression and persistence of PIT. Here we also wished to compare the sexes on their response to the water-associated cue, following water devaluation. Thus, using the rats in Experiment 2, we assessed the effects of prior water devaluation on cue-triggered increases in instrumental responding for water reward across the sexes. To this end, all rats were given a final PIT test session (42 min). Half of the male and half of the female rats (n = 8/sex) received free water access for 1 hour immediately before the start of this final PIT test session, while the other half (n = 8/sex) did not. We measured lever responses and water cup entries throughout the PIT session.

##### Statistical Analysis

Data were analysed using Graph Pad Prism. In Exps. 1 & 2, three-way mixed model ANOVA was used to analyse both active vs. inactive lever-pressing across instrumental conditioning sessions (Lever × Session × Sex; ‘Lever’ and ‘Session’ as within-subjects variables), and CS+ vs. CS- water cup entries across Pavlovian conditioning sessions (CS × Session × Sex; ‘CS’ and ‘Session’ as within-subjects variables). During PIT tests in Exp. 1, lever presses during CS+ vs. CS- were analysed using a three-way mixed model ANOVA (Lever × CS × Sex; ‘Lever’ and ‘Session’ as within-subjects variables), and the number of water cup entries and locomotor activity during CS+ vs. CS- presentations were analysed using a two-way mixed model ANOVA (CS × Sex; ‘CS’ as a within-subjects variable). During PIT tests in Exp. 2, a two-way repeated measures ANOVA was used to analyse LY379268 effects on lever pressing, water cup entries and locomotion (LY379268 × CS; all as within-subjects variables). A two-way mixed model ANOVA was used to analyse devaluation effects on lever pressing and water cup entries (Devaluation × CS; ‘CS’ as a within-subjects variable). When interaction and/or main effects were significant, Bonferroni-adjusted multiple post-hoc comparisons were used to analyse further effects. When Mauchly’s test of sphericity revealed a significant violation, the Greenhouse-Geisser correction was applied (for ε < 0.75). Across experiments, during PIT tests, we present lever pressing, water cup entries and locomotion rates during each CS (CS+ and CS-) presentations as elevation scores over baseline responding, given by the 2-min ITI period immediately prior to each 2-min CS presentation. Data are expressed as mean ± SEM. The α level was set at p ≤ 0.05.

## Results

### Exp. 1: Cue-triggered increases in incentive motivation for water reward in female and male rats

Rats received instrumental and then Pavlovian conditioning, before being tested for PIT (Fig. 1a). Rats increased their active lever pressing across instrumental training sessions (Fig. 1b; ANOVA; main effect of Session, F(8,240) = 18.58, p < 0.001). Rats also pressed more on the active versus inactive lever (main effect of Lever, F(0.6279, 18.84) = 42.6, p < 0.001, ε = 0.63), and this difference increased across sessions (Lever × Session interaction, F(4.844, 145.3) = 23.14, p < 0.001, ε = 0.61). This indicates that female and male rats effectively learned to lever press for water. Furthermore, males pressed more on the active lever than females did during instrumental training (Sex × Lever × Session interaction, F(8,240) = 1.46, p = 0.17; main effect of Sex, F(1, 30) = 15.25, p < 0.001; Sex × Lever, F(1, 30) = 22.51, p < 0.001; Sex × Session, F(8,240) = 1.66, p = 0.11), earning on average 35% more water than females did per session (note that males also weighed 31% more than females did; data not shown). Thus, all rats learned to lever press for water reward, and male rats responded more for water than female rats did, likely due to their larger size.

Rats next underwent Pavlovian conditioning. Fig. 1c shows that across Pavlovian conditioning sessions, rats increased their average rates of water cup entries during the first 10 sec of CS presentation (main effect of Session, F(7,210) = 14.34, p < 0.001). Rats also entered the water cup at a greater rate during CS+ compared to CS- presentations (Fig. 1c; main effect of CS, F(0.57,17.11) = 38.64, < 0.001, ε = 0.57), with this difference increasing across sessions (CS × Session interaction, F(5.075, 152.2) = 9.45, p < 0.001, ε = 0.72). This conditioned discrimination between CS+ and CS- was similar between female and male rats (main effect of Sex, F(1,30) = 0.43, p = 0.52; Sex × CS, F(1,30) = 1.30, p = 0.26; Sex × Session, F(7,210) = 0.88, p = 0.53; Sex × CS × Session interaction, F(7, 210) = 1.10, p = 0.36). Thus, across the sexes, rats learned the CS+/water contingency, and this Pavlovian conditioning was similar in female and male rats.

#### Cue-triggered increases in incentive motivation for water reward

During PIT tests, rats had access to both levers, but lever pressing was not reinforced by water or the CS+. Across CS types, rats pressed significantly more on the active than on the inactive lever, and did so when PIT tests were given both early (Day 1; Figs. 2a and c; main effect of Lever, F(1,30) = 17.77, p < 0.001) and late (Days 21/27; Fig. 2b and d; F(1,30) = 28.30, p < 0.001) after initial instrumental/Pavlovian conditioning. Thus, rats sought water during PIT testing and did so for up to 4 weeks after initial conditioning.

**Fig. 2.**
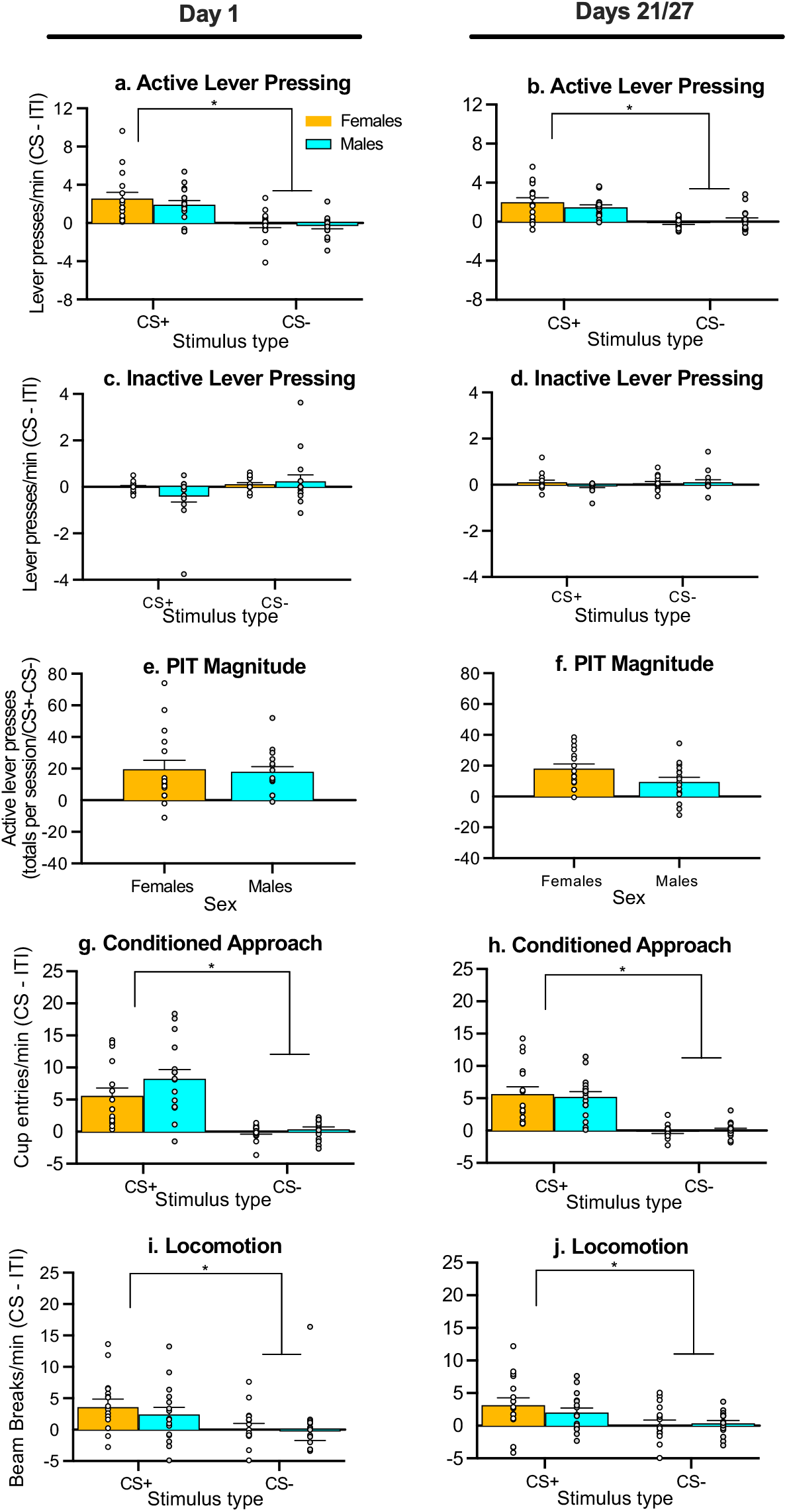
Female and male rats showed similar Pavlovian-to-Instrumental transfer (PIT). (a) Female and male rats pressed more on the previously water-associated lever when presented with the CS+ vs. the CS-, and this effect was observed both early, (b) and late after initial conditioning. (c-d) Across the sexes, CS+ and CS- presentations had similar effects on inactive lever presses. (e) PIT magnitude was similar between females and males tested for PIT early, (f) and late after initial conditioning. (g) Female and male rats showed significant cue-triggered discrimination of conditioned approach behaviour, visiting the water cup more often during CS+ compared to CS- presentations, and this effect was seen both early, (h) and late after initial conditioning. (i) Female and male rats showed similar increases in locomotion when the CS+ vs. CS- was presented, and this was the case both early (j) and late after initial conditioning. Data are means ± SEM (n = 16/sex). * *p* < 0.05. CS; conditioned stimulus. ITI; inter-trial interval.

Across the sexes, rats pressed more on the active lever during CS+ versus CS- presentations, and did so during tests given both early (Day 1; Figs. 2a and c; main effect of CS, F(1,30) = 19.62, p < 0.001; CS × Lever interaction, F(1,30) = 37.02, p < 0.001; active lever pressing during CS+ > CS-, p < 0.001) and late after initial instrumental/Pavlovian conditioning (Days 21-27; Figs. 2b and d; main effect of CS, F(1,30) = 24.95, p < 0.001; CS × Lever interaction, F(1,30) = 36.36, p < 0.001; active lever pressing during CS+ > CS-, p < 0.001). There were no sex differences in this effect (Day 1, Figs. 2a and c; main effect of Sex, F(1,30) = 1.07, p = 0.31; Sex × CS interaction, F(1,30) = 1.11, p = 0.30; Sex × Lever interaction, F(1,30) = 0.34, p = 0.57; Days 21-27, Figs. 2b and d; main effect of Sex, F(1,30) = 0.39, p = 0.54; Sex × CS interaction, F(1,30) = 2.18, p = 0.15; Sex × Lever interaction, F(1,30) = 0.05; All P’s > 0.05). Across the 2 test timepoints, females and males also showed similar PIT magnitude, (Figs. 2e-f; main effect of Sex, F(1,30) = 1.09, p = 0.31; Sex x Day interaction, F(1,30) = 1.77, p = 0.19). Thus, for up to 4 weeks after initial conditioning, presentation of the water-paired cue (CS+), but not the control cue (CS-) potentiated the operant pursuit of water reward, and this effect was comparable between female and male rats.

Rats entered the water cup more often during CS+ versus CS- presentations, and this was the case both early (Day 1; Fig. 2g; main effect of CS, F(1,60) = 48.18, p < 0.001) and later after initial instrumental/Pavlovian conditioning (Days 21-27; Fig. 2h; main effect of CS, F(1,60) = 58.23, p < 0.001). This conditioned approach response was similar between females and males (Fig. 2g; main effect of Sex, F(1,60) = 2.59, p = 0.11; Sex × CS interaction, F(1,60) = 1.26, p = 0.27; Fig. 2h; main effect of Sex, F(1,60) = 0.01, p = 0.91; Sex × CS interaction, F(1,60) = 0.25, p = 0.62). Thus, rats showed significant cue-triggered discrimination of conditioned approach to the site of water delivery for up to 4 weeks after initial conditioning, and this response was similar in female and male rats.

Finally, female and male rats significantly increased their locomotor activity when the CS+ versus CS- was presented, and this was observed at both testing time points (Day 1; Fig. 2i; main effect of CS, F(1,30) = 7.88, p = 0.01; Days 21-27; Fig. 2j; main effect of CS, F(1,60) = 8.40, p = 0.01). This locomotor response was similar across the sexes (Day 1; Fig. 2i; main effect of Sex, F(1,30) = 0.36, p = 0.55; Sex × CS interaction, F(1,30) = 0.13, p = 0.72; Days 21-27; Fig. 2j; main effect of Sex, F(1,60) = 0.25, p = 0.62; Sex × CS interaction, F(1,60) = 0.79, p = 0.38). Thus, rats showed enhanced locomotor activity during CS+ vs. CS- presentations for up to 4 weeks after initial conditioning, and this effect was comparable in female and male rats.

### Exp. 2: Effects of systemic injection of an mGlu2/3 receptor agonist on cue-triggered increases in incentive motivation for water reward

Rats first received instrumental and Pavlovian conditioning, then, two weeks later, we assessed the effects of systemic injections of the mGlu2/3 agonist, LY379268 on cue-elicited increases in instrumental responding for water reward (i.e., PIT; Fig. 3a). Across instrumental training sessions, rats increased their rates of lever pressing (Fig. 3b; main effect of Session, F(8,240) = 29.55, p < 0.001), and they pressed more on the active vs. inactive lever across sessions (main effect of Lever, F(0.6174, 18.52) = 257.5, p < 0.001, ε = 0.62; Lever × Session interaction, F(4.821, 144.6) = 38.69, p < 0.001, ε = 0.60). Compared to females, males pressed more on the active lever, and also showed a greater increase in active lever pressing across sessions (Fig. 3b; Sex × Lever × Session interaction, F(8,240) = 4.59, p < 0.001; main effect of Sex, F(1,30) = 15.53, p < 0.001; Sex × Lever, F(1,30) = 13.28, p = 0.001; Sex × Session, F(8,240) = 3.46, p < 0.001; active lever pressing, males > females, p < 0.001). Consequently, males earned more water than female rats did, specifically during the first two days of responding under a variable interval schedule of water reinforcement (Fig. 3c; main effect of Sex, F(1,30) = 11.63, p = 0.002; Sex × Session interaction, F(8,240) =2.16, p = 0.03; during VI10-1 and VI10-2 sessions, males > females, p = 0.002 and p < 0.001, respectively. No other comparisons where statistically significant). Female and male rats earned progressively fewer water deliveries/session over time (Fig. 3c; main effect of Session, F(1.85, 55.62) = 56.11, p < 0.001, ε = 0.23), due to the increasing VI schedules of reinforcement used. Thus, as in Experiment 1, all rats learned to lever press for water reward, and while male rats lever pressed more for water than female rats did, the two sexes earned a similar amount of water on the last 6 out of 8 instrumental conditioning sessions.

**Fig. 3.**
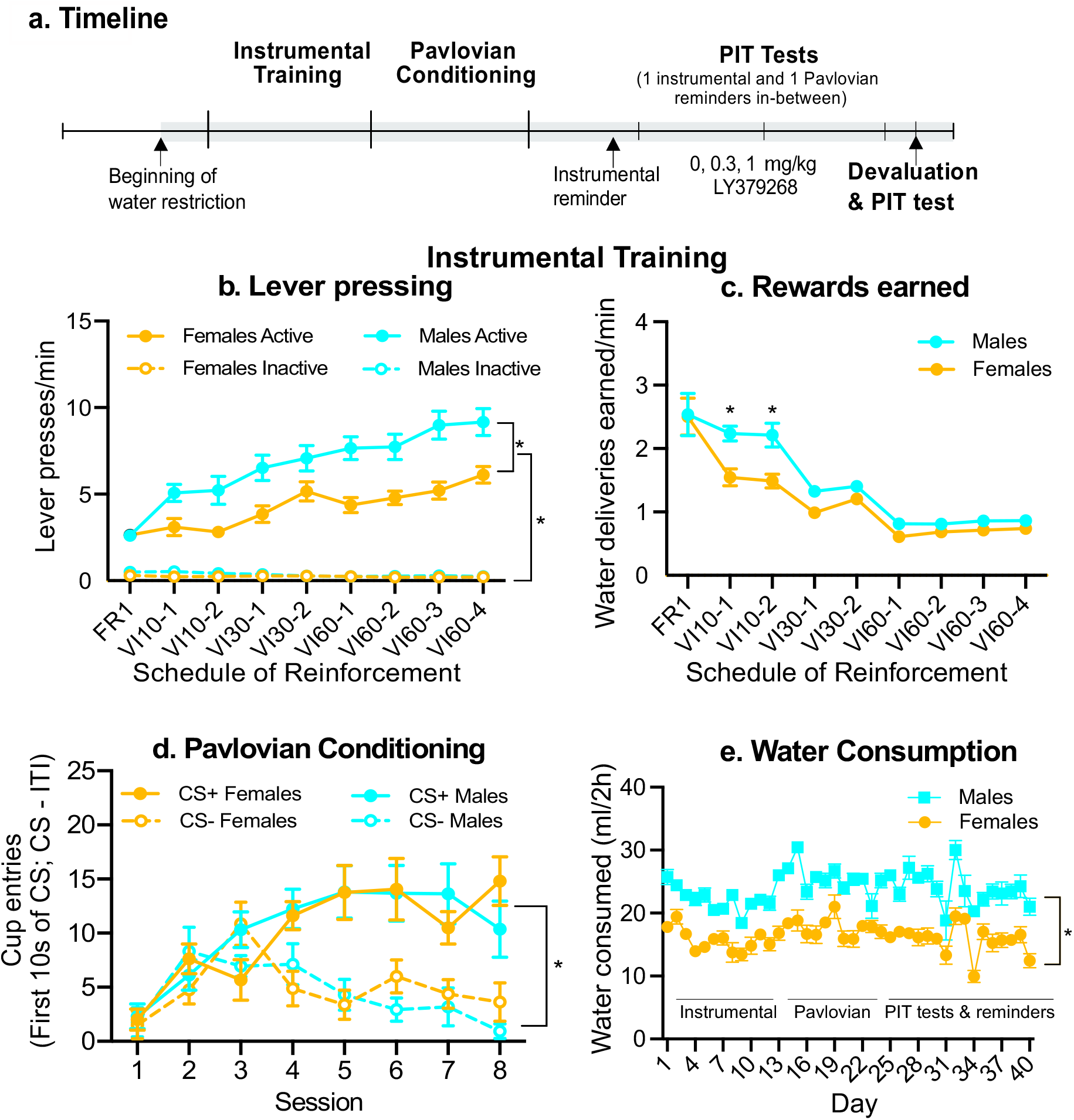
Timeline and acquisition of instrumental and Pavlovian conditioning in Experiment 2. (a) Female and male rats underwent instrumental and Pavlovian conditioning. Two weeks after the last conditioning session, we assessed the influence of subcutaneous injections of the mGlu_2/3_ receptor agonist, LY379268 (0, 0.3 and 1 mg/kg) on cue-triggered increases in incentive motivation for water, using Pavlovian-to-instrumental transfer (PIT) procedures. One day after the last LY379268 injections test, we assessed the influence of reward devaluation on the expression of PIT. (b) The rate of active lever presses increased over the daily 40-min instrumental training sessions, and males pressed more often on the active lever than females did. (c) During instrumental training, males also earned more water than females did. (d) The rate of water cup entries during the first 10 sec of each CS presentation increased over the daily 44-min Pavlovian conditioning sessions, and there were no sex differences in this effect. (e) Males consumed more water than females did during the 2-h daily water access periods. Data are means ± SEM (n = 16/sex). \**p* < 0.05. FR; fixed ratio. VI; variable interval. CS; conditioned stimulus. ITI; inter-trial interval.

Over the course of Pavlovian conditioning, rats increased their average rates of water cup entries during the first 10 sec of CS presentation (Fig. 3d; main effect of Session, F(7,210) = 7.69, p < 0.001). Rats also visited the water cup more frequently during CS+ compared to CS- presentations (main effect of CS, F(0.5645, 16.93) = 53.96, p < 0.001, ε = 0.56), and this conditioned discrimination increased across sessions (CS × Session interaction, F(4.205, 126.1) = 8.74, p < 0.001, ε = 0.60). There were no sex differences in this response (main effect of Sex, F(1, 30) = 0.01, p = 0.93; Sex × CS, F(1,30) = 0.28, p = 0.60; Sex × Session, F(7,210) = 0.97, p = 0.45; Sex × CS × Session interaction, F(7, 210)= 1.74, p = 0.10). Thus, rats showed CS+ triggered conditioned approach to the site of water delivery, and females and males did so to a comparable degree.

As Fig. 3e shows, in Experiment 2, we measured daily water consumption in females and males during the 2-h/day access periods. Male rats consumed more water than female rats did (main effect of Sex, F(1,14) = 68.77, p < 0.001; Sex × Day interaction, F(39, 546) = 1.92, p = 0.001).

#### Effects of systemic LY379268 on lever pressing

Rats received systemic injections of LY379268 (0, 0.3 and 1 mg/kg, s.c.) 30 min before being tested for PIT. As in Exp. 1, females and males showed similar behavioural responses during baseline PIT testing (i.e., when injected with vehicle; All P’s > 0.05). The sexes also responded similarly to LY379268 during PIT tests (All P’s > 0.05). Thus, all PIT data were collapsed across the sexes.

An initial analysis across all PIT test sessions showed that rats pressed more often on the previously water-associated (active) lever during CS+ water cue vs. CS- presentations (Fig. 4a; main effect of CS, F(1,179) = 41.95, p < 0.001). In contrast, rates of inactive lever pressing were similar during CS+ and CS- (Fig. 4a; p = 0.65). Thus, CS+ presentation selectively increased pressing on the previously water-associated lever, without increasing lever-pressing behaviour indiscriminately.

**Fig. 4.**
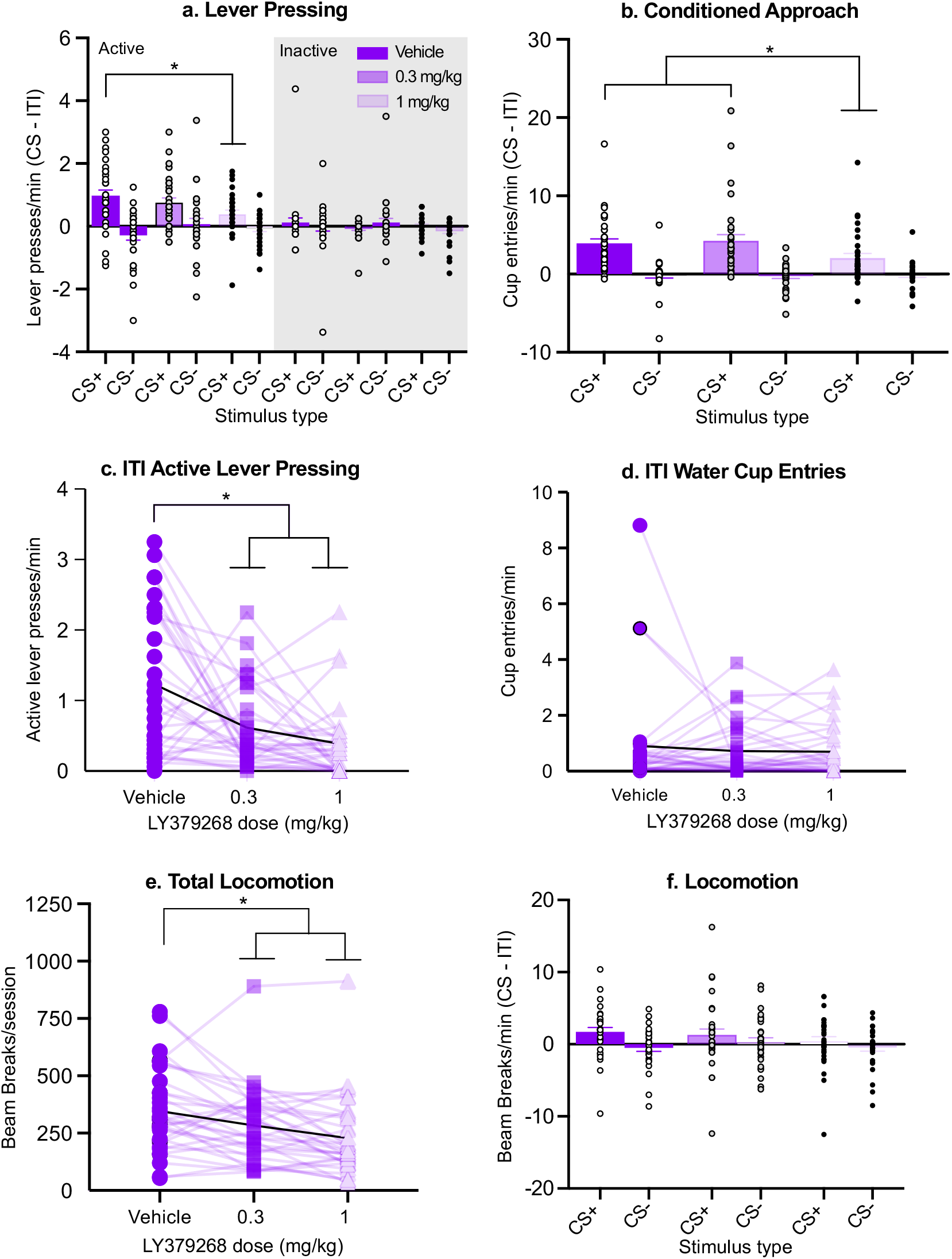
Systemic injection of the mGlu_2/3_ receptor agonist, LY379268 attenuated cue-triggered increases in both instrumental responding for water reward and conditioned approach behaviours to the site of water delivery in male and female rats. (a) During Pavlovian-to-instrumental transfer (PIT) tests, CS+ presentations invigorated lever pressing for water reward, and LY379268 (1 mg/kg) reduced this effect. LY379268 had no effect on inactive lever presses. (b) During PIT testing, the rate of water cup entries was significantly higher during CS+ versus CS- presentations, and LY379268 (1 mg/kg). reduced this Pavlovian conditioned approach behaviour. (c) During inter-trial intervals (ITI), LY379268 suppressed active lever presses, (d) but had no effect on water cup entries. (e) Similarly, LY379268 reduced total locomotor activity, (f) but not locomotor activity during CS presentations. There were no sex differences in any of these responses. In Panels (c), (d), and (e), the black curve represents group averages. Bar graphs present data as means ± SEM (N = 32; half female, half male). * *p* < 0.05. CS; conditioned stimulus.

The effects of LY379268 on lever pressing during PIT tests depended upon CS type, such that LY379268 dose-dependently attenuated active lever pressing during CS+, but not during CS- presentations (Fig. 4a, active lever; CS × LY379268 interaction, F(2,179) = 3.53, p = 0.03; during CS+ presentations, 1 mg/kg LY379268 < Vehicle; p = 0.026. No other comparisons were statistically significant). LY379268 did not influence responding on the inactive lever (Fig. 4a, inactive lever; All P’s > 0.05). This suggests that the mGlu2/3 receptor agonist selectively decreased lever pressing for water reward, without influencing lever-pressing behaviour indiscriminately. However, rates of inactive lever pressing were already low under baseline (vehicle) conditions, indicating a potential floor effect. This being said, the data indicate that activation of mGlu2/3 receptors attenuated CS+ triggered potentiation of water-seeking behaviour, suggesting suppressed cue-triggered increases in incentive motivation for water.

Fig. 4b shows that analysis of water cup approach data during PIT tests, a measure that reflects the conditioned anticipation of water reward. Rats visited the water cup more often during CS+ vs. CS- presentations (Fig. 4b; main effect of CS, F(1,31) = 38.99, p < 0.001). Thus, the rats showed cue-triggered discrimination of approach to the site of water delivery. LY379268 dose-dependently reduced water cup entries, and did so specifically during CS+ presentations (main effect of LY379268, F(2,62) = 3.10, p = 0.052; LY379268 × CS interaction, F(2,62) = 4.07, p = 0.02; during CS+, Vehicle > 1 mg, p = 0.01; 0.3 mg > 1 mg, p = 0.002; during CS-, all P’s > 0.05. Note however that rats rarely entered the water cup during CS- presentations, indicating a potential floor effect). Thus, the CS+, but not the CS- increased the frequency of visits to the water cup, and activating mGlu2/3 receptors reduced this CS+-induced conditioned approach behaviour.

During PIT tests, LY379268 suppressed some general motor behaviours and left others unchanged. Fig. 4c shows that compared to vehicle, LY379268 reduced rates of active lever pressing during intervals between CS+/CS- presentations (inter-trial intervals or ITI; F(2,62) = 12.94, p < 0.001; Vehicle > 0.3 mg, p = 0.002; Vehicle > 1 mg, p < 0.001). However, Fig. 4d shows that LY379268 did not influence the rates of water cup entries during the ITI (F(1.267, 39.28) = 0.29, p = 0.64, ε = 0.63). LY379268 also reduced total locomotor activity counts (Fig. 4e; F(2,62) = 11.65, p < 0.001; Vehicle > 1 mg, p < 0.001; Vehicle > 0.3 mg, p = 0.04), but not locomotor activity during CS presentations (Fig. 4f; Main effect of LY379268, F(2,62) = 1.37, p = 0.26; LY379268 × CS interaction, F(2,62) = 0.63, p = 0.54). In summary, LY379268 attenuated lever pressing for water reward during inter-trial intervals and total locomotion levels, but LY379268 left both water cup entries during inter-trial intervals and locomotion during CS presentations unaffected.

#### Effects of prior water devaluation on lever pressing

Finally, after quantifying LY379268 effects on cue-triggered responding for water reward, we gave all rats a final PIT test (without LY379268 injections). Immediately before this test, half of the rats of each sex received water access for 1 h, to examine the influence of reward devaluation on PIT. Despite prior reward devaluation, rats pressed more often on the previously water-associated (active) lever during CS+ versus CS-presentation (Fig. 5a; main effect of CS, F(2,60) = 17.49, p < 0.001). However, reward devaluation produced an overall reduction of the incentive motivation for water, seen as reduced responding on the previously water-associated lever across CS+, CS- and ITI periods (Fig. 5a; main effect of Devaluation, F(1,30) = 12.77, p = 0.001; Devaluation × CS interaction, F(2,60) = 2.33, p = 0.11).

**Fig. 5.**
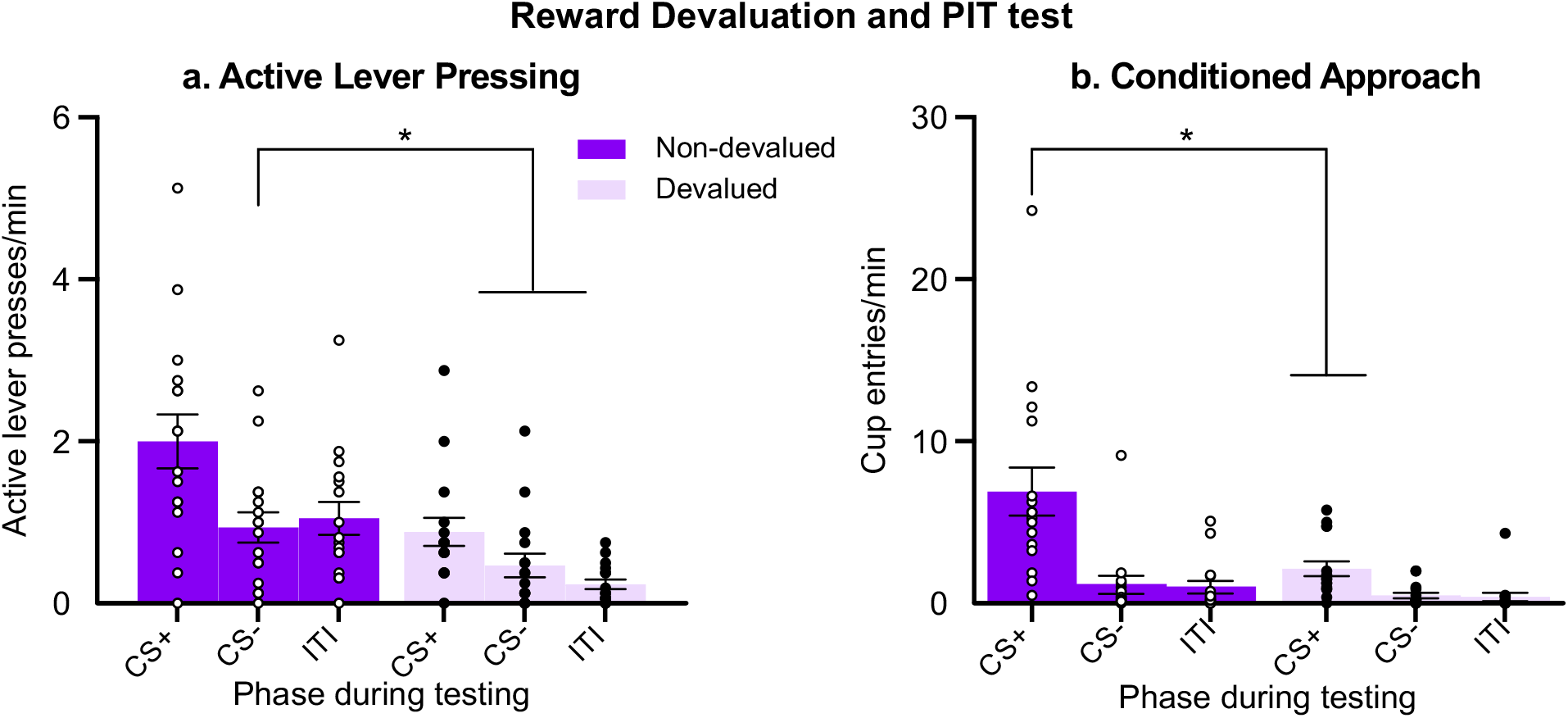
Water reward devaluation attenuated both lever pressing for this reward and cue-triggered approach behaviours to the site of water delivery in male and female rats. (a) During tests for Pavlovian-to-instrumental transfer (PIT), CS+ presentations increased ongoing lever pressing for water reward, and prior water devaluation reduced this instrumental response. (b) During PIT testing, rats entered the water cup significantly more often during CS+ versus CS- presentations, and prior devaluation of water reward decreased this Pavlovian conditioned approach response. Data are means ± SEM (n = 16/condition, half female, half male). * *p* < 0.05. CS; conditioned stimulus.

The rats visited the water cup more frequently during CS+ presentations vs. during CS- presentations or the ITI (Fig. 5b; main effect of CS, F(1.110, 33.29) = 26.03, p < 0.001, ε = 0.55), and devaluation reduced this cue-triggered discrimination of conditioned approach response (Fig. 5b; main effect of Devaluation, F(1,30) = 8.09, p = 0.01; Devaluation × CS interaction, F(2,60) = 7.81, p < 0.001; CS+ Non-devaluation > CS+ Devaluation, p = 0.02. No other comparisons were statistically significant). Thus, letting rats drink water before testing (i.e., devaluing water reward) reduces both responding on the previously water-associated lever and CS+ triggered increases in conditioned anticipation of water reward. This suggests that under our conditions, CS+ evoked potentiation of responding for water reward remains a deliberate, goal-directed behaviour, rather than a habitual response that relies exclusively on past instrumental/Pavlovian experience.

## Discussion

Here we compared the sexes on cue-triggered potentiation of water reward-seeking behaviours. We report three key findings. First, female and male rats show comparable expression and persistence of PIT, responding more for water during CS+ versus CS- presentations, and doing so both early (Day 1) and later (Days 21-27) after initial conditioning. This indicates that the sexes show similar cue-triggered potentiation of incentive motivation for water reward. Second, in female and male rats, water devaluation reduced both water-seeking behaviour and cue-triggered conditioned approach to the water port. Finally, activation of mGlu2/3 receptors with a selective agonist suppressed cue-triggered increases in both water reward seeking and water anticipation, and this effect was comparable in females and males.

### Biological sex influences performance during instrumental training, but not during Pavlovian conditioning or PIT tests

Male rats showed higher rates of lever-pressing for water during instrumental training relative to females, and they also earned more water during initial instrumental conditioning sessions. However, this effect was short-lived, such that on the last 6 of 8 instrumental sessions, males and females earned similar amounts of water. The schedule of reinforcement became progressively more demanding across training sessions. It is possible that male rats earned more water on the first few sessions because response requirements were lower during these earlier sessions, and the males’ larger bodies require more water (see Fig. 3e). Females and males showed similar performance during Pavlovian conditioning. The acquisition of CS+ triggered conditioned approach behaviour was similar between the sexes. Finally, even though males consume more water and can also self-administer more water during instrumental sessions, females and males showed similar CS+-triggered increases in both water-seeking actions and conditioned approach behaviours during PIT testing. We also found no sex differences in the persistence of these effects. Thus, for up to 4 weeks after initial conditioning, females and males showed comparable CS+ evoked potentiation of water-seeking and conditioned approach behaviours.

Our findings are consistent with work showing similar cue-induced reinstatement of cocaine-seeking in female and male rats (Feltenstein et al., 2011). Note also that Fuchs, Evans, Mehta, Case, and See (2005) found no effect of estrous phase on cue-induced reinstatement of extinguished reward seeking. To our knowledge, there are no published accounts of how hormonal fluctuations might influence PIT in females. This being said, in a meta-analysis of neuroscience-related traits, Becker, Prendergast, and Liang (2016) found that while there can be instances where females are more variable than males, there is an ‘overall absence of sex differences in variability across diverse traits of interests to neuroscientists’, such that ‘inclusion of intact females, without regard to estrous cycle, and intact males is a valid approach to learn about females in neuroscience research’.

### Activation of mGlu2/3 receptors attenuates cue-triggered increases in incentive motivation for reward

We found that in female and male rats, systemic administration of an mGlu2/3 receptor agonist (LY379268) impairs the ability of a water-associated cue to increase both water-seeking actions and conditioned anticipation of water reward. This concords with studies showing that mGlu2/3 receptor activity mediates cue-triggered appetitive behaviours, such as cue-induced reinstatement of reward-seeking behaviour (Backstrom & Hyytia, 2005; Baptista et al., 2004; Bossert, Poles, et al., 2006; Lu et al., 2007; Uejima et al., 2007; Zhao et al., 2006).

Here, LY379268 decreased cue-induced potentiation of reward seeking in a pure conditioned incentive motivation paradigm. The PIT procedure we used allowed us to selectively assess the influence of mGlu2/3 receptor activity on incentive motivation, because PIT measures the ability of a cue to trigger motivation for a reward, without the confounding influences of either primary or secondary reinforcement (Balleine, 1994; Dickinson & Balleine, 1994; Dickinson & Dawson, 1987; Rescorla & Solomon, 1967; Wyvell & Berridge, 2000). As such, the effects of mGlu2/3 receptor activation likely do not involve any suppression of the primary reinforcing value of water, because rats were tested under extinction conditions, when lever pressing no longer produced water. Our water consumption data also show that on PIT test days, when rats received the mGlu2/3 receptor agonist, they consumed as much water in their home cages after testing as they had on preceding days (Fig. 3e). This further supports the idea that administration of an mGlu2/3 receptor agonist does not significantly change the reinforcing properties of water.

Our results cannot be due to the ability of mGlu2/3 receptor activity to mediate conditioned reinforcement. Pavlovian cues can act as conditioned reinforcers, strengthening preceding instrumental responses. To act as a conditioned reinforcer, a reward cue must be presented contingently after an instrumental response. However, the PIT procedure we used precluded any such reinforcement contingency, as the cue was presented independently of lever-pressing behaviour. Thus, the results suggest that pharmacological activation of mGlu2/3 receptors decreases the pure conditioned incentive impact of reward cues, without necessarily influencing primary or secondary reinforcement. As such, mGlu2/3 receptor activity could have suppressed cue effects on reward-seeking behaviour by decreasing the incentive salience of the cue-triggered neural representation of water reward (see Wyvell & Berridge, 2000).

The neurobiological mechanisms involved in the ability of mGlu2/3 receptor activation to attenuate cue-induced incentive motivation for reward remain to be studied. However, the observed attenuation of cue-triggered incentive motivation following mGlu2/3 activation is consistent with a reduction in synaptic glutamate release. We did not measure extracellular glutamate concentrations here. However, mGlu2/3 receptor activity provides autoregulatory, negative feedback to decrease stimulated glutamate release (Conn & Pin, 1997; Imre, 2007; Schoepp, 2001). Several studies suggest that cue-triggered increases in incentive motivation require synaptic glutamate in regions such as the amygdala (Derman, Bass, & Ferrario, 2020; Feltenstein & See, 2007; Garceau et al., 2021; Khoo, LeCocq, Deyab, & Chaudhri, 2019; Lu et al., 2007; Malvaez et al., 2015) and the nucleus accumbens (Derman & Ferrario, 2018). Thus, our findings extend work showing that synaptic glutamate transmission is necessary for cue-triggered increases in incentive motivation.

We found that systemic LY379268 attenuated some motor measures, but not others. LY379268 reduced total locomotor activity and lever pressing behaviour during inter-trial intervals (i.e., in the absence of the CS+). Previous studies using male rats have also noted motor effects with systemic LY379268 (Allain et al., 2017; Backstrom & Hyytia, 2005; Kufahl, Martin-Fardon, & Weiss, 2011), but at doses higher than used here (≥ 2 mg/kg). For instance, Backstrom and Hyytia (2005) found that at 5 mg/kg, LY379268 decreased spontaneous locomotor activity in male rats. Other studies (Allain et al., 2017; Cannady, Grondin, Fisher, Hodge, & Besheer, 2011; Kufahl et al., 2011) found that the doses used here (0.3 or 1 mg/kg) did not significantly impair motor behaviour (but 2 and 3 mg/kg did). There are differences in the species used in these previous experiments compared to the present one. Cannady et al. (2011) used Long-Evans rats, others used Wistar rats (Allain et al., 2017; Kufahl et al., 2011), while here we used Sprague-Dawley rats. The apparent discrepancies between our findings and these previous experiments may arise from inherent species differences in sensitivity to the motor-impairing effects of LY379268. Sensitivity to the motor-impairing effects of LY379268 is also brain region-dependent. For example, the amygdala is relatively insensitive to LY379268’s motor effects (e.g., at 6 μg/hemisphere), while lower doses (3 μg/hemipshere) injected into the nucleus accumbens can reduce locomotion (Besheer et al., 2010; Cannady et al., 2011). This suggests that systemic LY379268 may evoke some motor-suppressing effects in part by diffusing to sites of action in the nucleus accumbens. Indeed, systemic LY379268 can reduce activity in neuronal projections from the nucleus accumbens to the globus pallidus (Cannady et al., 2011), a structure that coordinates locomotor function (Mogenson, Swanson, & Wu, 1983). Thus, in the present work, motor suppression could have contributed at least partially to the effects of LY379268 on CS-evoked increases in incentive motivation.

However, it is unlikely that the effects of the mGlu2/3 receptor agonist on PIT are driven by motor actions alone. First, LY379268 completely spared motor behaviours including locomotor activity during CS presentations and water cup entries during inter-trial intervals. Second, during inter-trial intervals, water cup entries occurred at a similar rate as active lever pressing did (compare Figs. 4c and d). However, LY379268 injection only suppressed the latter behavioural response. Thus, a simple motor-suppression explanation of behaviour cannot account for the effects of LY379268 during PIT testing. Second, other work shows that LY379268 can decrease cue-triggered potentiation of incentive motivation for reward without having non-specific motor effects. For example, injecting LY379268 into the BLA reduces both cue-evoked potentiation of incentive motivation for water reward and CS-induced anticipation of reward, without affecting other motor behaviours such as locomotion, and active lever pressing or water cup entries during inter-trial intervals (Garceau et al., 2021). Of course, the effects of systemic injections of LY379268 could be different. Whatever the case may be, future work can determine the extent to which motor suppression contributes to LY379268’s effects on the behavioural response to reward cues using different mGlu2/3 receptor agonists and/or different doses of LY379268.

### Prior reward devaluation attenuates cue-triggered increases in incentive motivation for reward

When rats were allowed to freely drink water for 1 h before PIT testing, thus devaluing the water reward, they showed reduced responding on the water-associated lever and reduced conditioned approach to the water cup. There were no significant differences between females and males in this effect. Previous studies on the influence of devaluation on cue-triggered increases in incentive motivation report mixed results, possibly due to the different procedures used. In rats, devaluation by satiation suppresses general, but not specific PIT (Corbit, Janak, & Balleine, 2007; Holland, 2004). While devaluation by conditioned taste aversion (i.e., pairing the reward with illness induced by lithium chloride) had no effect on PIT in a single-lever paradigm (Holland, 2004), satiation abolished PIT under the same test conditions (Aitken, Greenfield, & Wassum, 2016; Dailey, Moran, Holland, & Johnson, 2016). The latter finding is consistent with our results, as we used a single-outcome PIT paradigm and observed that devaluation by satiation suppressed both reward-associated lever responding and cue-triggered increases in conditioned approach. The palatability of the primary reward can also influence the response to devaluation by satiation. For instance, Kendig, Cheung, Raymond, and Corbit (2016) showed that devaluation by satiation is less effective in reducing food-seeking behaviour in the presence of contexts or discrete cues paired with junk food, compared to when rats were in the presence of contexts and cues paired with less palatable, regular chow. Here, we used plain water as the US, which is less reinforcing after satiation because water’s reinforcing properties depend upon the level of thirst. This could explain the suppressive effects of devaluation we found on water-associated lever responding. Thus, water-seeking responses in our paradigm were sensitive to devaluation, suggesting this instrumental responding was likely goal-directed, and not habitual (Balleine & Dickinson, 1998).

## Conclusions

In conclusion, a water-associated cue increases both conditioned anticipation of water reward and water-seeking actions, and does so to a similar extent in female and male rats. Water devaluation by satiation also reduced water-associated lever responding and cue-triggered conditioned anticipation of water reward in both sexes. Finally, our results confirm and extend previous findings, showing that in both sexes, activating mGlu2/3 receptors with systemic injections of LY379268 decreases both reward-seeking actions when the reward is not available (i.e., under extinction conditions), and cue-triggered increases in conditioned approach to the site of water delivery. These effects were seen at an LY379268 dose that also suppressed some motor measures. However, motor deficits are unlikely to fully account for LY379268’s effects on PIT, because the agonist left other motor measures unchanged. Thus, we hypothesize that signaling via mGlu2/3 receptors mediates the motivating influence of appetitive cues, thereby influencing the ability of such cues to goad reward-seeking actions when the associated reward is not immediately available.

## Acknowledgements

This work was supported by the National Science and Engineering Research Council of Canada (grant 355923) and the Canada Foundation for Innovation to ANS (grant 24326). ANS holds a salary award from the Fonds de la Recherche du Québec-Santé (Grant #28988). CG was supported by a Master’s scholarship (B1X) from the Fonds de recherche du Québec - Nature et technologies (Grant #275340). The authors gratefully acknowledge Rifka (Rebecca) C. Derman, Carrie R. Ferrario and Cameron Nobile for generously sharing equipment, computer programs and advice in setting up the behavioural tasks used in this work. The authors declare no conflicts of interest.

